# Construction of a three-component regulatory network of transcribed ultraconserved regions for the identification of prognostic biomarkers in gastric cancer

**DOI:** 10.1101/2022.08.12.503817

**Authors:** Anis Khalafiyan, Modjtaba Emadi-Baygi, Markus Wolfien, Ali Salehzadeh-Yazdi, Parvaneh Nikpour

**Author notes:** **Correspondence should be addressed to:** Parvaneh Nikpour (PhD), Department of Genetics and Molecular Biology, Faculty of Medicine, Isfahan University of Medical Sciences, Isfahan, Iran.

## Abstract

Although altered expression and functional roles of the transcribed ultraconserved regions (T-UCRs) in the pathophysiology of neoplasms has already been investigated, relevance of the functions for T-UCRs in gastric cancer (GC) is still the subject of inquiry. In the current study, The Cancer Genome Atlas Stomach Adenocarcinoma (TCGA-STAD) dataset was used as a resource for the RNA-sequencing data. Differential expression analysis was conducted using DESeq2. Interactions between T-UCRs, miRNAs, and mRNAs were combined into a three-component network. The Survival package was utilized to identify survival-related differentially-expressed T-UCRs (DET-UCRs). Using an in-house cohort of GC tissues, expression of two DET-UCRs was experimentally verified. Thirty-four T-UCRs were dysregulated in TCGA-STAD tumoral samples compared to non-tumoral counterparts. The network was composed of 34 DET-UCRs, 275 miRNAs and 796 mRNAs nodes. Five T-UCRs were significantly correlated with the overall survival. While no expression of uc.232 was observed in our in-house cohort of GC tissues, uc.343 showed an increased expression in gastric tumoral tissues. The constructed three-component regulatory network of T-UCRs in GC presents a comprehensive understanding of the underlying gene expression regulation processes involved in tumor development and can serve as a basis to investigate potential prognostic biomarkers and therapeutic targets.

**Simple summary:** Gastric cancer (GC) is one of the most common cancers in the world and is considered as a highly heterogeneous disease based on subtypes and genetic alterations. GC is mostly detected in the advanced stages, hence, identifying diagnostic and prognostic biomarkers is of urgent need. Transcribed ultraconserved regions (T-UCRs) are a type of long non-coding RNAs which are linked to human carcinogenesis. Their mechanisms of action and the factors regulating their expression in cancers are poorly understood. In the current study, by applying a systems biology approach and constructing a regulatory network, we have presented a T-UCR as a potential diagnostic biomarker. Additionally, five T-UCRs with significant correlation with patients’ overall survival were found, which can be potentially used as prognostic biomarkers in future.

## Background

Gastric cancer (GC), with more than one million new cases diagnosed globally each year, is the sixth most common cancer and the third main cause of cancer-related mortality in the world [1]. Gastric cancer has been histopathologically divided into two subtypes of intestinal and diffuse [2], however it is regarded as a highly heterogeneous disease with respect to architecture and growth, cell differentiation, and molecular pathogenesis [3].

As in the majority of cases, GC is diagnosed in an advanced stage; conventional treatment options are not effective leading to a poor prognosis with 10–12 months median overall survival (OS) [4, 5]. Therefore, establishing novel, sensitive, and specific tools to identify diagnostic and prognostic biomarkers for GC are urgently demanded.

Progression in transcriptome analyses has disclosed that ∼98% of human DNA is transcribed into RNAs, which do not translate to proteins and are entitled as non-coding RNAs (ncRNAs) [6-8]. Long noncoding RNAs (lncRNAs) are classified as ncRNAs with a length of at least 200 nucleotides. They are critical regulatory factors in various biological and pathological processes, like differentiation [9], immune response [10], metabolism [11], and cancer development and progression [12-14].

Transcribed-ultraconserved regions (T-UCRs) are a new type of lncRNAs that was found in 2004 after bioinformatics analyses of mouse, rat, and human genomes. They are 481 genomic elements (range: 200–779 bp) encoded by a subset of DNA ultraconserved regions that show 100 percent homology between orthologous loci in the human, mouse, and rat genomes [15, 16]. Expression of T-UCRs demonstrate both ubiquitous and tissue-specific patterns [16]. Only 9% of T-UCRs are bi-directionally transcribed, with most showing transcription from one strand [16]. Primary mechanisms regulating the expression of T-UCRs include interactions with microRNAs (miRNAs) and hypermethylation of CpG island promoters [16, 17]. T-UCRs are mostly located in the fragile sites and cancer-associated genomic regions (CAGRs) [16] acting as oncogenes or tumor suppressors [17, 18]. Current information suggest that T-UCRs can provide well-suited candidates as diagnostic and prognostic biomarkers.

In 2016, Goto et al. conducted qRT-PCR analysis on 14 T-UCR regions on GC that had previously shown reduced expression in prostate cancer tissue samples, reporting that eight T-UCRs including uc.359 + A, uc.282 +, uc.261 +, uc.252 + A, uc.249 +, uc.244 +, uc.158 + A and uc.118 + A had a decreased expression in gastric tumoral compared to control tissues. Moreover, uc.416 + A showed significant overexpression in tumoral GC samples. Further analysis revealed that uc.416 + A is associated with tumor cell growth supporting its role as an oncogene in GC [18]. Furthermore, overexpression of uc.63+ and downregulation of uc.160+ in GC tumoral tissues have been reported in other studies as well [19-21].

Thus far, no comprehensive network construction and analysis has been performed on the expression of all T-UCRs along with miRNAs, and mRNAs in gastric cancer. In this study, we aimed to estimate the overall expression and potential functions of all T-UCRs in gastric cancer, as well as their association with prognosis.

## Materials and Methods

### Ethics Statement

This study protocol was approved by the Ethics Committee of Isfahan University of Medical Sciences (IR.MUI.MED.REC.1398.443) and was in accordance with the Declaration of Helsinki. Furthermore, approval for the controlled-access data (RNA-seq binary alignment map (BAM) files) was collected from dbGaP [22].

### Identification of differentially-expressed T-UCRs

Sequence data in the BAM file format (373 files of stomach adenoma and adenocarcinoma (STAD) containing 343 tumor and 30 non-tumor samples) as well as clinical data from The Cancer Genome Atlas (TCGA) database were downloaded from the GDC data portal (https://portal.gdc.cancer.gov). We linked the BAM and clinical files via patient identifiers. Of these, 343 files were derived from tumor samples and the rest were taken from normal samples. The entire list of human-rat-mouse ultraconserved sequences [15] was downloaded from https://users.soe.ucsc.edu/~jill/ultra.html. CrossMap [23] in Ensemble was used to convert genomic coordinates of T-UCRs to the hg38 human genome assembly. For quantification of the T-UCR expression, featureCounts version 1.5.0 [24] was applied to obtain raw count values of T-UCRs in each sample. Differential expression analysis was then performed using DESeq2 to identify potential deregulation of T-UCRs. The fold change (FC) of individual T-UCR levels was computed and differentially-expressed T-UCRs (DET-UCRs) were considered significant with |log2 FC| > 1 and an adjusted p-value < 0.05.

### Construction of a T-UCR-miRNA-mRNA network

After identifying differentially-expressed T-UCRs and to construct a three-component regulatory network, sequences of T-UCRs were obtained from supplementary data by Bejerano et al. [15] to identify the potential binding sites for miRNAs in T-UCRs sequence. In particular, these sequences were converted into reverse-complementary RNA using an online software: Reverse Complement from bioinformatics portal of The Sequence Manipulation Suite (http://www.bioinformatics.org/sms/rev_comp.html). Sequences of miRNAs were retrieved from the online database miRBase release 19 [25]. Human miRNAs with”high confidence”, highly conserved identities in different mammalian species, were selected. Then, we applied the RNAhybrid software (v 2.1, Bielefeld, Germany) available online at http://bibiserv.techfak.unibielefeld.de/rnahybrid/ [25, 26] as a target prediction tool to determine putative miRNA target sites in T-UCR sequences [25]. To choose RNA duplexes between T-UCRs and miRNAs, we considered high stringency conditions, such as constraint of seed matching to 2–7 nucleotides and a p-value < 0.05 [26]. Then, to predict the interaction between miRNAs and mRNAs, we utilized the online database: miRWalk3.0 (http://mirwalk.umm.uni-heidelberg.de/) [27]. The miRNA-mRNA interactions, which were common between three databases of miRDB, miRTarBase, and TargetScan, were finally selected. A resulting network was constructed and visualized with Cytoscape (version 3.7.2). In the network, the molecular components are considered as “nodes”, while their interactions are referred to as “edges”. A node with a large degree of centrality is considered as a hub.

### Functional enrichment analysis

To identify the most significant enriched pathways in the network, an enrichment analysis was carried out for mRNAs using DAVID (version 6.8) (Database for Annotation, Visualization, and Integrated Discovery, http://david.abcc.ncifcrf.gov/) [28]. Gene Ontology (GO) and Kyoto Encyclopedia of Genes and Genomes (KEGG) with a p-value < 0.05 were selected for functional enrichment analysis. The ggplot2 R software was used to visualize the results [29].

### Protein-protein interaction (PPI) network analysis

To investigate the function of mRNAs also at the protein level within a connected network, we utilized the STRING database [30]. A PPI network with the combined scores greater than 0.9 and hiding disconnected nodes was then constructed. Cytoscape (version 3.7.2) was used to compose the PPI network [31]. The KEGG pathways retrieved from highly-connected proteins were also explored using STRING interaction data.

### Survival analysis

To identify survival-related differentially-expressed T-UCRs, we used Kaplan-Meier survival curves. Log rank p-value < 0.05 and Hazard Ratio (HR)≠1 were considered as statistically significant thresholds. Based on the cut-off point, the median T-UCR’s expression levels of the TCGA samples were grouped as high or low. Kaplan-Meier survival curves were constructed using the survival package (https://CRAN.R-project.org/package=survival) in R (version 3.6.3).

### Clinical specimens

A total of 64 tumoral and adjacent non-tumoral tissues were obtained from patients with GC for the validation of the identified biomarkers. The specimens were collected by the Iran National Tumor Bank, funded by the Cancer Institute of Tehran University, for Cancer Research as described previously [32, 33]. Prior to participating, all patients gave written informed consent to the Iran Tumoral Bank.

### RNA extraction and reverse transcription polymerase chain reaction (RT-PCR)

QIAzol® lysis reagent (Qiagen, USA) was used to extract total RNA from tissues according to the manufacturer’s instructions. The integrity of the RNA was determined using a 1% agarose gel electrophoresis, and the concentration of the RNA was determined using a Nanodrop instrument (NanolytiK, Duesseldorf, Germany). DNase I treatment was conducted to prepare DNA-free RNA prior to RT-PCR using the DNase kit (Thermo Scientific, Germany). Random hexamer primers and an M-MLV reverse transcriptase were used to synthesize complementary DNA (cDNA) from DNase-treated RNA (Yekta Tajhiz Azma, Iran). On a Magnetic Induction Cycle (Mic) instrument (Bio Molecular system, Australia), qRT-PCR was performed using RealQ Plus 2x Master Mix Green High ROX (Ampliqon, Denmark) and specific primers for two candidate T-UCRs and *ACTB* as a reference gene (Table 1). The RT-PCR conditions were as follows: a primary denaturation at 95°C for 15 minutes, 40 cycles of denaturation at 95°C for 15 seconds, annealing at 60°C for 30 seconds, and extension at 72°C for 30 seconds. As a positive control, genomic DNA isolated from human blood was used in all experiments. All reactions were carried out at least in duplicate. The relative gene expression of the candidate T-UCRs were calculated using the 2^−ΔΔCt^ method [34]. Furthermore, two PCR products were sequenced with an Applied Biosystems 3100 Genetic Analyzer to verify that the T-UCRs are being specifically amplified.

**Table 1:**
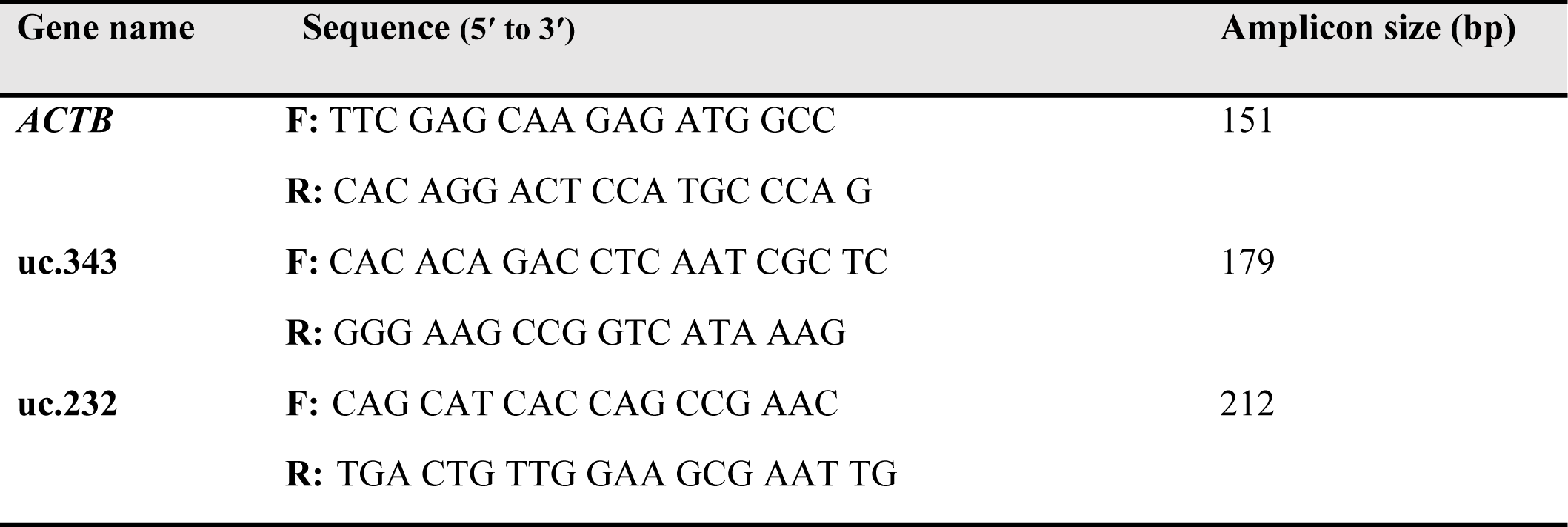
Sequences of real-time quantitative PCR primers in this study

### Statistical analysis

The normal distribution of the samples was determined using the Shapiro-Wilk test in SPSS software, version 16.0 (SPSS Inc., Chicago, USA). The statistical significance was then determined using the Student’s t/Mann-Whitney U test in Prism 7.0 (Graph-Pad Prism Software Inc., San Diego, CA). The qRT-PCR results were presented as mean ± standard error of the mean (SEM). Statistical significance was defined as a p-value of less than 0.05.

## Results

### Differential expression analysis of T-UCRs

The workflow for the integrated bioinformatics analysis and experimental validation is shown in Figure 1.

**Figure 1.**
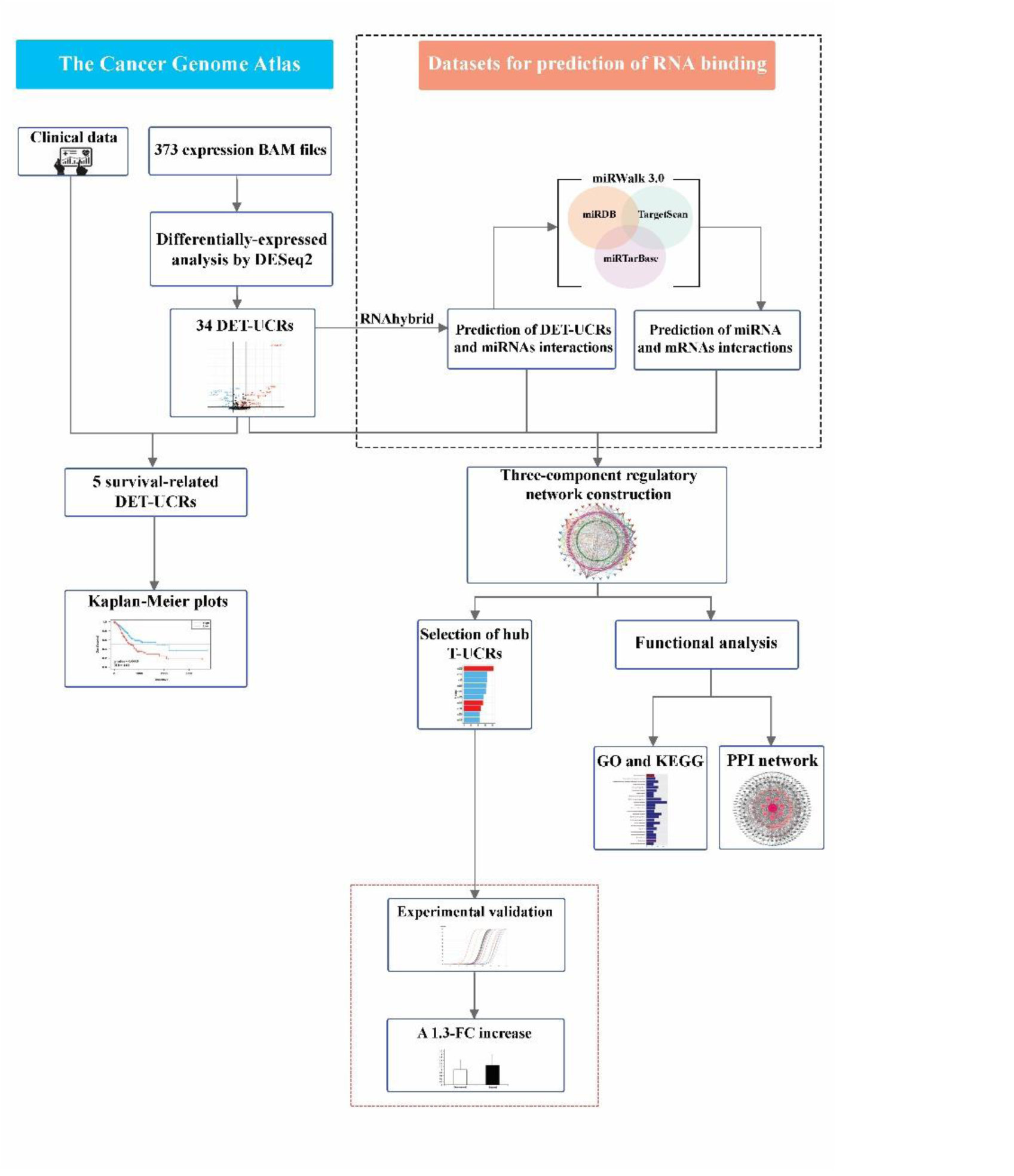
The workflow of the study design and analysis. The black dashed box represents the steps for predicting miRNAs interacting with T-UCRs and mRNAs interacting with these miRNAs. The red dashed box shows the experimental validation of candidate biomarkers on our own collected cohort of GC tissues. BAM: Binary Alignment Map, DET-UCRs: differentially-expressed T-UCR, FC: fold change, GO: gene ontology, KEGG: Kyoto Encyclopedia of Genes and Genomes, PPI: protein-protein interaction

To determine the differentially-expressed T-UCRs (DET-UCRs) in GC, we compared the expression of 481 known T-UCRs in TCGA tumor samples compared to non-tumor samples. The respective analysis of RNA-seq data identified 34 DET-UCRs at a statistically significant level (adjusted p value < 0.05). Of these 34 T-UCRs showing differential, 14 were upregulated and 20 were downregulated in gastric tumoral tissues compared to normal ones (Figure 2).

**Figure 2.**
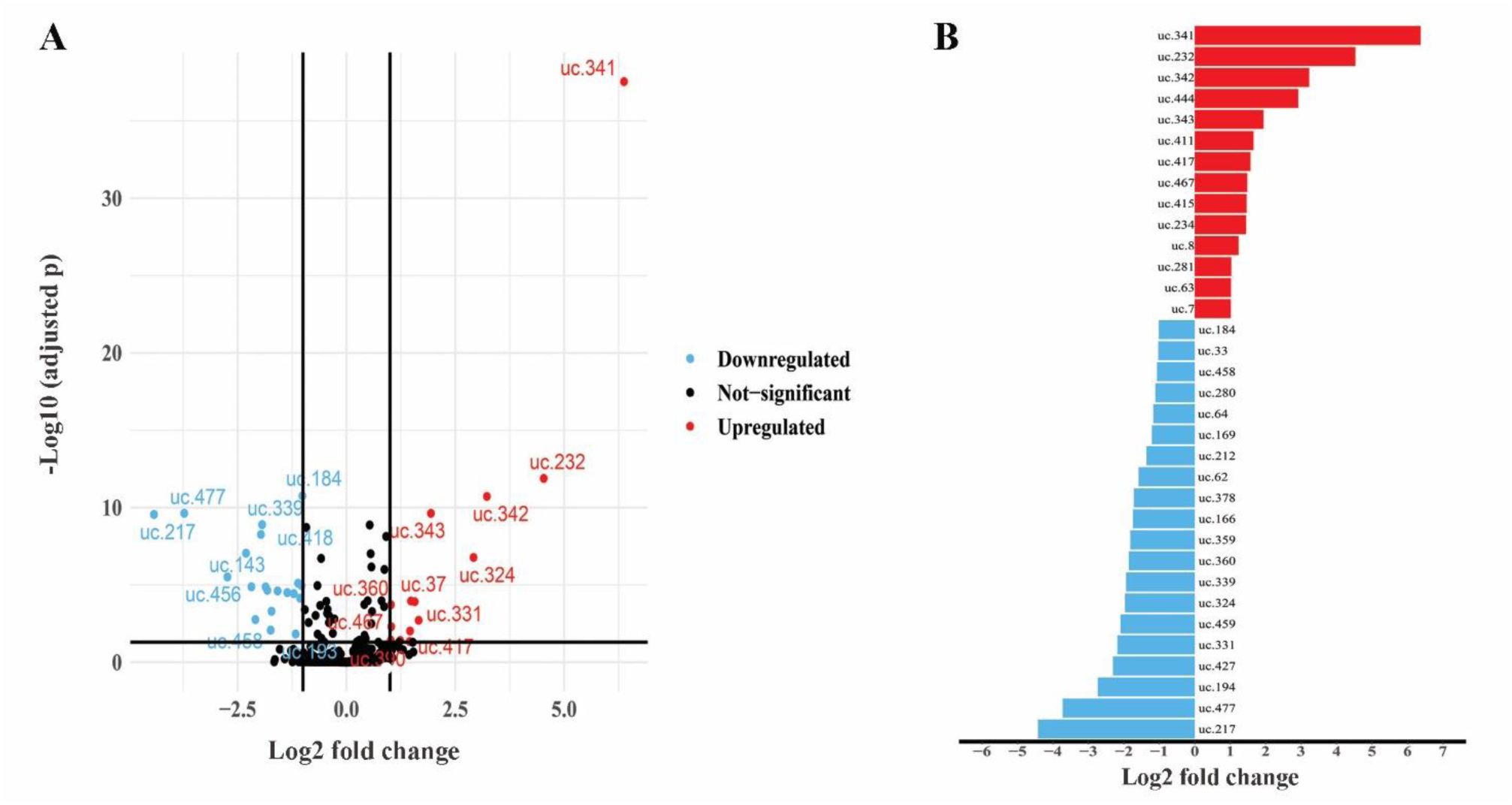
Significantly differentially expressed T-UCRs in gastric cancer. Volcano plot (A) and bar plot (B) of differentially-expressed T-UCRs between TCGA gastric tumor and non-tumor tissues. The blue dots/bars represent downregulated T-UCRs, the red dots/bars represent upregulated T-UCRs, and the black dots represent T-UCRs that were not significantly differentially expressed T-UCRs with |Log_2_ fold change (FC)| > 1 and an adjusted p-value < 0.05 were considered as significantly differentially expressed T-UCRs (DET-UCRs). TCGA: the cancer genome atlas, T-UCRs: transcribed ultraconserved regions

### Regulatory network of differentially-expressed T-UCRs in GC

A three-component regulatory network in GC composed of DET-UCRs, their associated miRNAs, and target mRNAs was constructed. As shown in Figure 3A, the three-component regulatory network consists of 34 DET-UCRs, 275 miRNAs, and 796 mRNAs nodes, as well as 1,284 edges (Supplementary file 1). In Figure 3B, the top ten highly connected T-UCRs (hubs) are ranked according to the degree centrality retrieved from the CytoHubba plugin. Moreover, the network has been made available on the Network Data Exchange ([three component network](https://www.ndexbio.org/#/network/404a7bb1-cc91-11ec-b397-0ac135e8bacf?accesskey=74406b2038d11866bf3aefcc7a7d7946ab61187806f3e198876e1d67d6056891)), a database and online community for sharing and collaborative development of network models, which can be easily imported into the Cytoscape for individual exploration [35].

**Figure 3.**
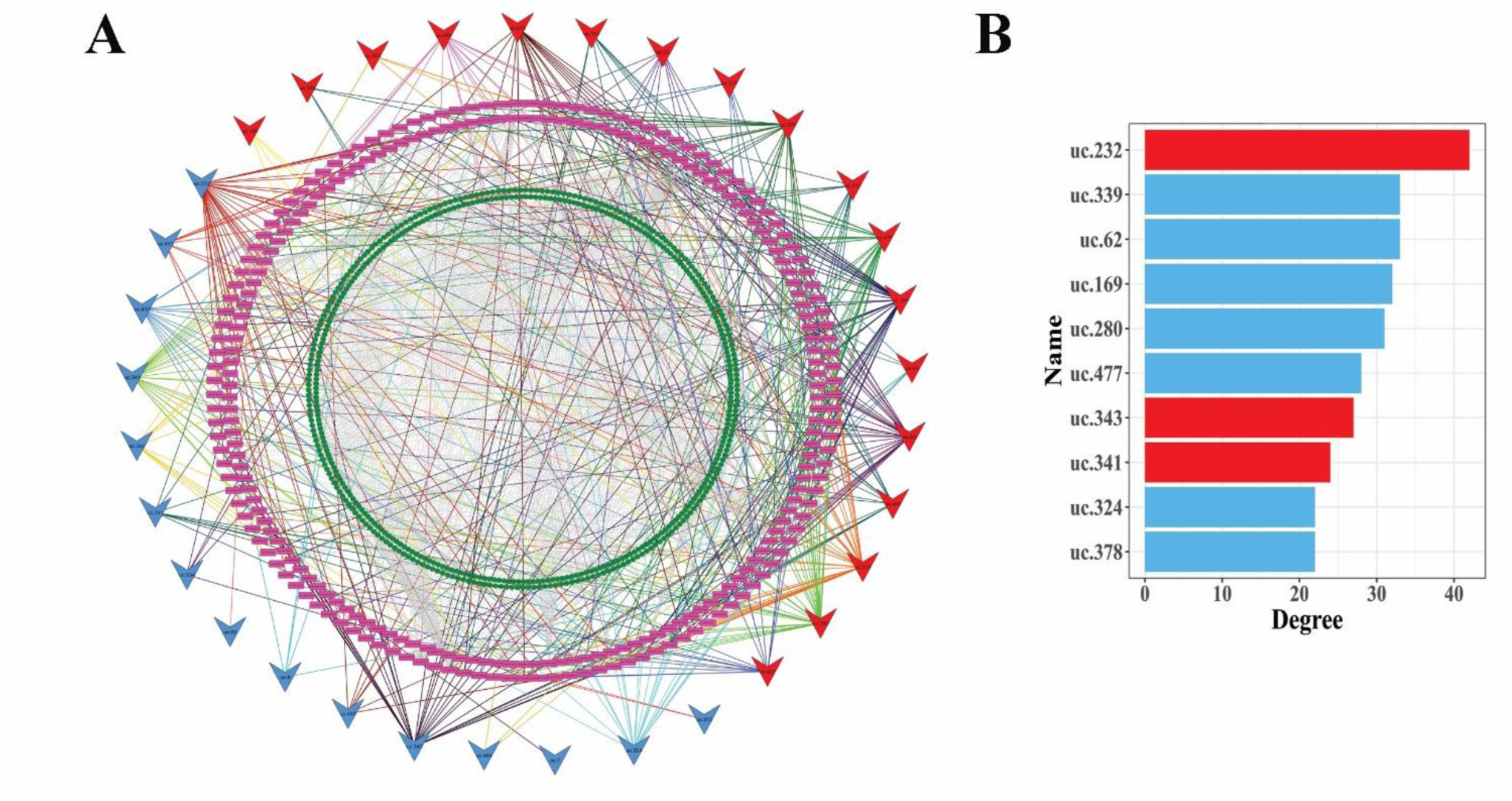
The three-component regulatory network. Red and blue V shapes represent upregulated and downregulated T-UCRs, respectively. Purple rectangles represent miRNAs and green circles represent mRNAs. Colored lines show predicted interactions between miRNA and T-UCR and grey dashes indicate interactions between miRNA and mRNA based on miRWalk3.0 (A). The top ten T-UCRs are ordered by network degree centrality. Upregulated and downregulated T-UCRs are represented by the red and blue bars, respectively (B). T-UCRs: transcribed ultraconserved regions

### Functional implication of network mRNAs

In order to predict the roles of T-UCRs in GC, we investigated the potential functional implications of the 796 mRNAs in the network, by functional annotations of GO and KEGG of these mRNAs (Supplementary file 2). The top 25 GO biological processes and molecular functions and top 25 KEGG pathways of mRNAs, based on the p-values, were selected for biological and functional analyses. The enriched GO biological processes of mRNAs were significantly involved in processes, such as “transcription”, “regulation of transcription”, “signal transduction”, “regulation of cell proliferation”, “regulation of apoptotic process”, and “cell cycle” (Figure 4A). GO molecular functions of mRNAs, which are presented in Figure 4B, include “protein binding”, “metal ion binding”, and “DNA binding”. The analysis of KEGG pathways showed that mRNAs were significantly enriched in pathways, such as “pathway in cancer”, “microRNAs in cancer”, and “PI3K-AKT signaling pathway” (Figure 4C).

**Figure 4.**
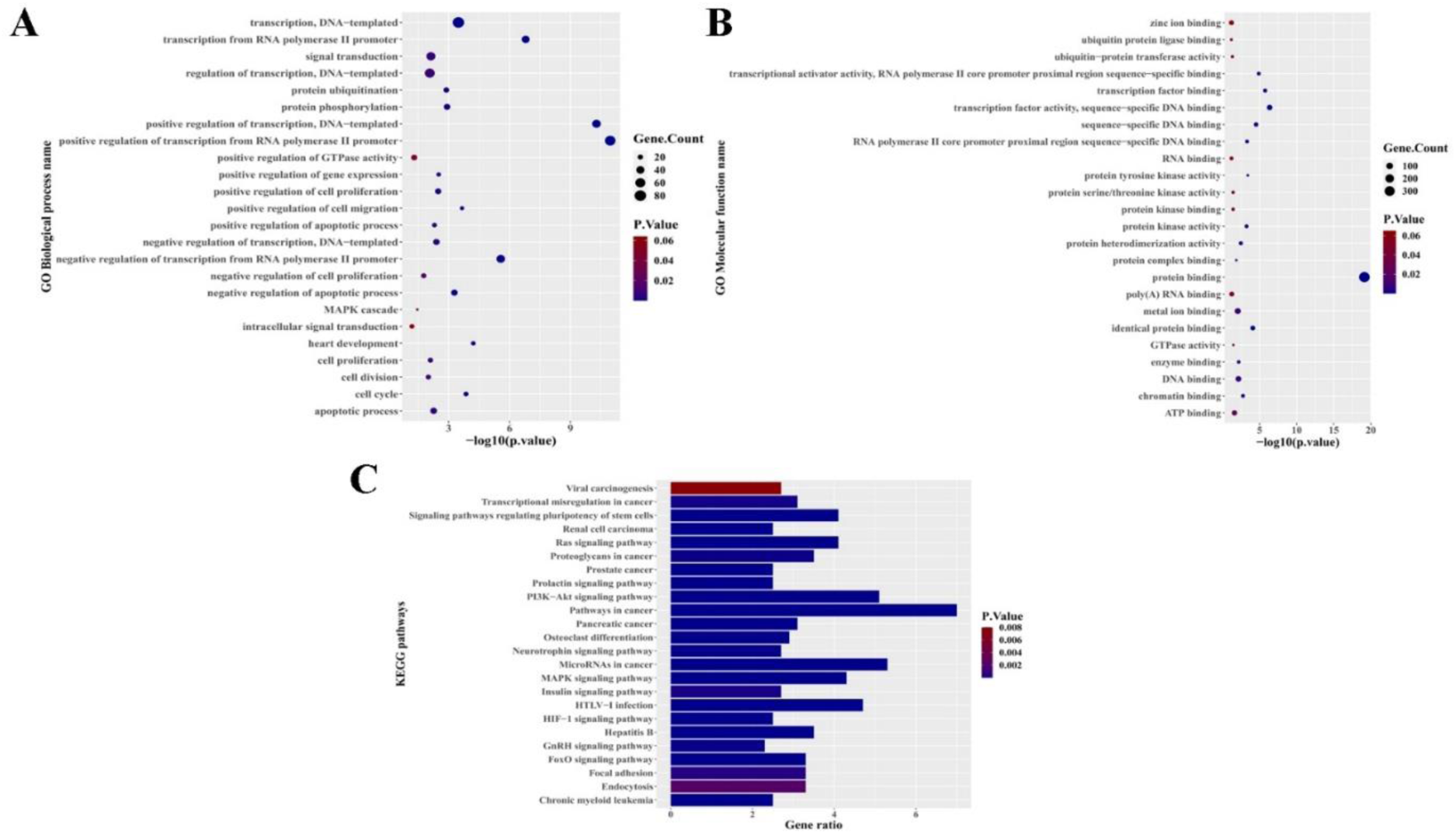
The enrichment analysis of GO and KEGG pathway for mRNAs in the network. Top 25 GO biological process terms (A), molecular functions (B), and KEGG pathways of mRNAs in the network (C). P-value < 0.05 was considered as statistically significant. Number of genes ascribed to a pathway within a specific list of differentially-expressed genes is shown by the Gene Ratio. GO: gene ontology, KEGG: Kyoto Encyclopedia of Genes and Genomes

### Functional implication of network mRNAs using protein-protein interaction analysis

By selecting the combined scores > 0.9 and hiding the disconnected nodes, the PPI network consisted of 198 nodes and 540 edges (Figure 5A). The top 20 predominant nodes with the highest degree centralities are shown in Figure 5B. Results of STRING analysis showed that these highly connected proteins have strong associations with prolactin signaling pathway, growth hormone synthesis, secretion and action, EGFR tyrosine kinase inhibitor resistance, signaling pathways regulating pluripotency of stem cells, and pathways in cancer (Supplementary file 3).

**Figure 5.**
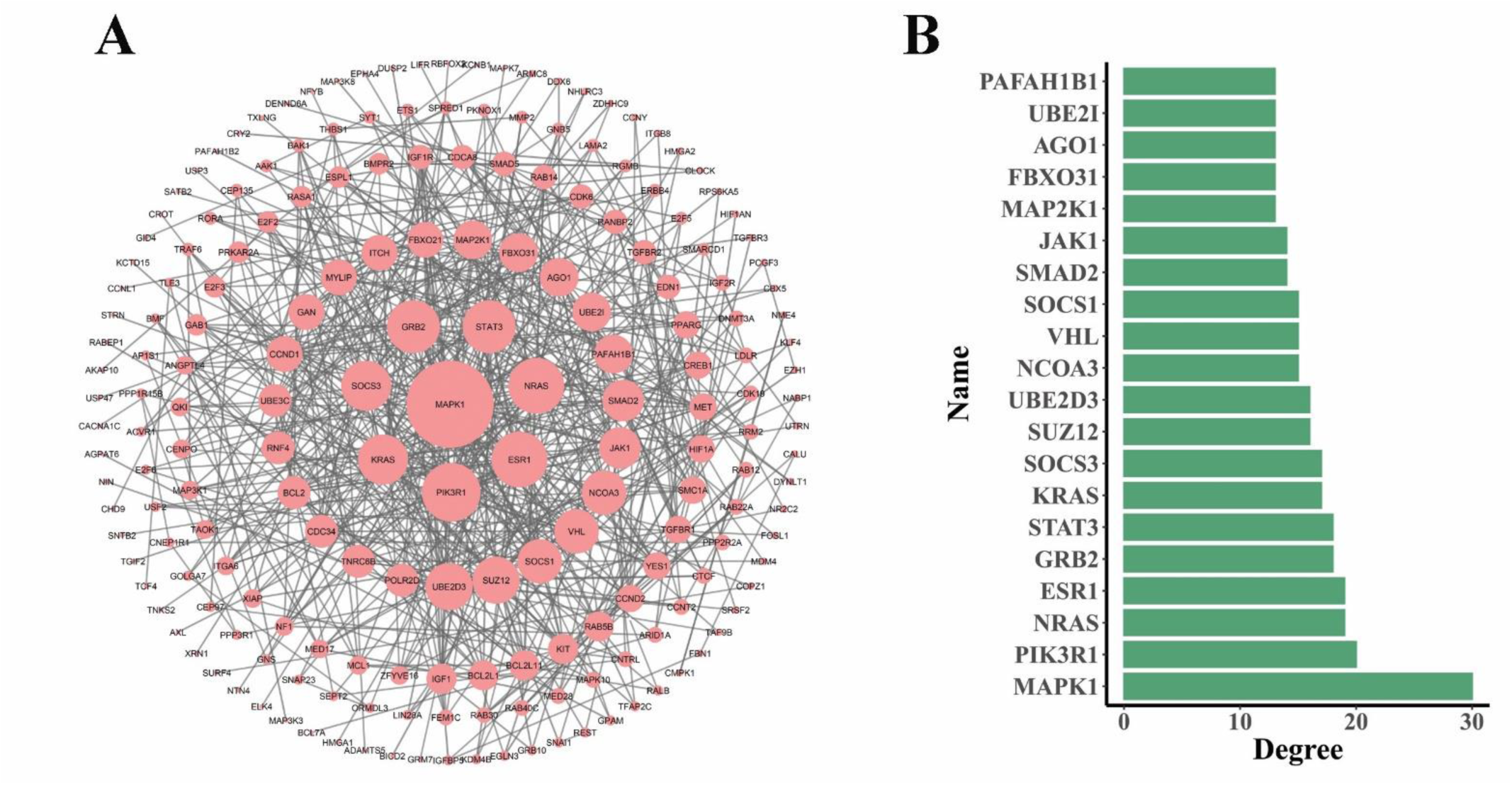
The protein-protein interaction (PPI) network of mRNAs. The nodes represent proteins (with a combined score of >0.9), and their size is proportional to their degree centrality (A). The diagram represents the top 20 predominant nodes with the highest degree centralities (B).

### Survival-related DET-UCRs

Univariate survival analysis was conducted to identify differentially expressed T-UCRs (DET-UCRs), which are associated with overall survival. A total of five DET-UCRs was revealed as survival-related T-UCRs in STAD based on log-rank p-value < 0.05 and HR ≠ 1. STAD patients with higher expression of uc.169 and uc.360 and lower expression of uc.8, uc.342, and uc.417 had significantly smaller survival rates (Figure 6).

**Figure 6.**
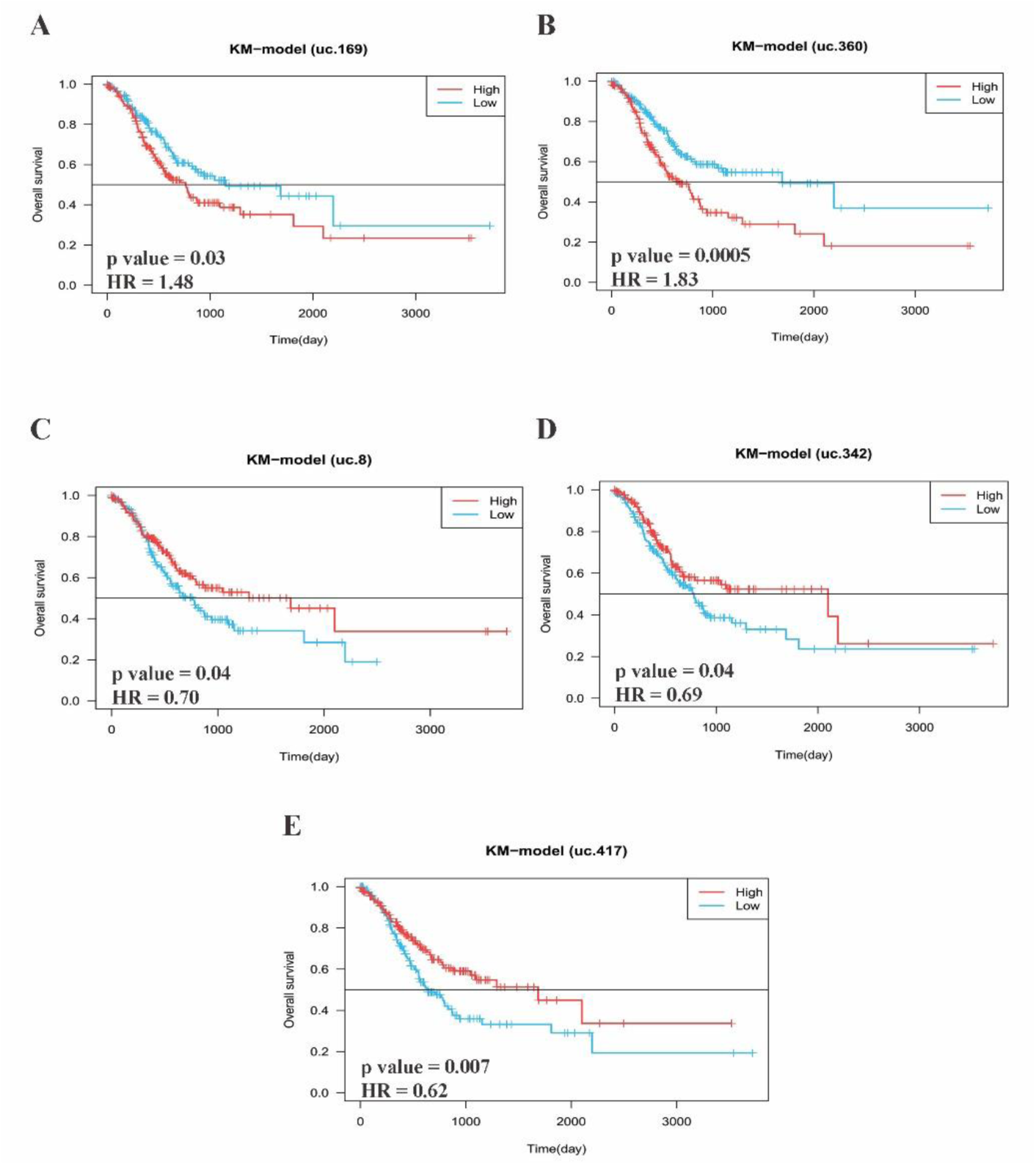
Kaplan-Meier survival curves of DET-UCRs. Based on a log-rank p-value < 0.05 and HR ≠ 1, five differentially-expressed T-UCRs were shown to be linked with overall survival. Patients with higher expression of uc.169 (A) and uc.360 (B) and lower expression of uc.8 (C), uc.342 (D) and uc.417 (E) had significantly poorer survival rates. The cutoff value for splitting STAD patients into two groups was determined by the median expression value of each T-UCR: red and blue lines indicate high and low expression of the T-UCRs, respectively. DET-UCRs: differentially-expressed T-UCRs, HR: hazard ratio

### Experimental validation of two candidate DET-UCRs

According to the degree centrality criteria in the constructed regulatory network, as well as showing upregulated patterns (Figure 2B), two candidate DET-UCRs, namely uc.232 and uc.343, were selected for further experimental evaluation. Prior to the quantitative analysis, optimization processes were carried out on a genomic DNA as a positive control for both ordinary and real-time RT-PCR reactions, using specific primers for uc.232 and uc.343. Agarose gel electrophoresis of the PCR products revealed single bands of the predicted sizes for the amplified uc.232 (212 bp) and uc.343 (179 bp) segments, respectively (Supplementary file 4). Real-time PCR analysis of gene expression revealed a unique melting curve without primer dimers (data not shown), which was further confirmed by agarose gel separation and staining. Two PCR products were also sequenced to demonstrate specific amplification of the transcripts. Database comparisons using nucleotide BLAST (https://blast.ncbi.nlm.nih.gov/Blast.cgi) revealed that the amplified PCR products were 100% identical to the uc.232 and uc.343 transcripts (Supplementary file 5). After optimizing the real-time PCR, the expression of uc.232 in tumoral and adjacent non-tumoral stomach tissues was investigated using real-time RT-PCR. Results showed that while expression of uc.232 in the gDNA sample was easily detectable, it was undetectable in all examined gastric tissues. Compared to non-tumoral gastric tissues, an increased expression of uc.343 was observed in tumoral gastric tissues, this increase, however, was not statistically significant (Figure 7).

**Figure 7:**
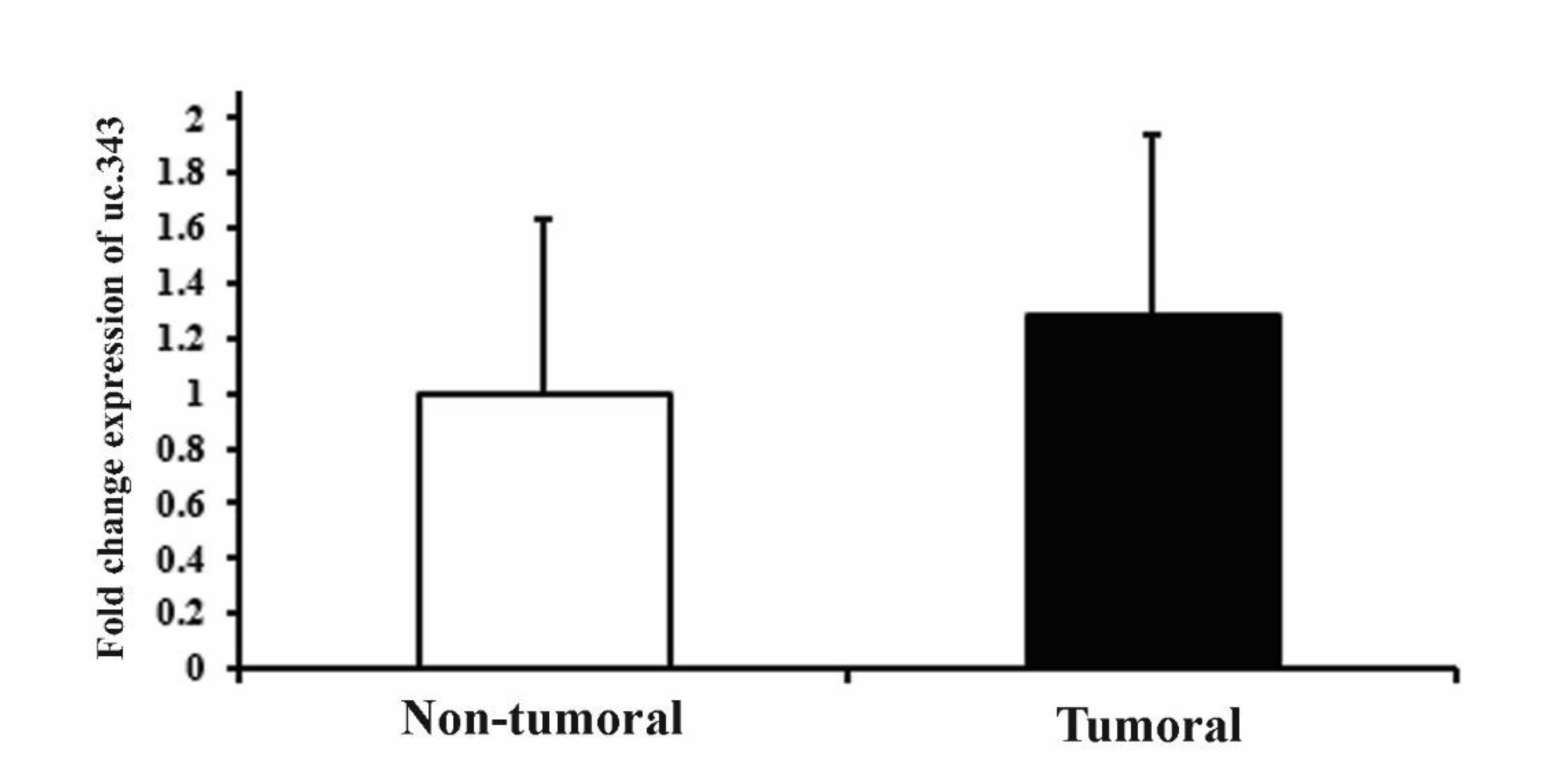
The expression level of uc.343 in tumoral and non-tumoral gastric tissue samples. Data are presented as mean ± standard error of the mean (SEM).

## Discussion

Gastric cancer is the world’s sixth most frequent cancer [1]. The incidence and mortality of GC have decreased considerably because of great advances in surgical procedure, chemotherapy, and targeted therapy during the last 50 years. As most patients with GC are without symptoms for years, they are more likely to be diagnosed in advanced stages. This could also be owing to the absence of appropriate diagnostic biomarkers for early detection of GC. For these reasons, studying gene expression profiles of GC can be an asset to identify and propose efficient diagnostic and prognostic biomarker candidates and predict response to treatments.

In the last few years, noncoding RNAs have emerged as important players in development of tumors [36], which includes its various classes (e.g., miRNAs and T-UCRs). In the pathogenesis of most cancers investigated, ncRNAs contribute an additional layer to the molecular architecture of human tumors [16, 37]. T-UCRs have been implicated in tumorigenesis according to recent research [38]. Furthermore, researchers have reported that T-UCRs can affect gene expression through direct binding to mRNA or miRNAs [39]. T-UCRs and their downstream targets can thus build intricate monitoring networks that are linked to the occurrence and progression of many malignancies.

In this study, we comprehensively examined the expression profiles of 481 T-UCRs in GC from the TCGA cohort. We identified 34 significantly differentially-expressed T-UCRs between GC tumors and normal samples that resulted in a disease-specific expression pattern. There are previous studies, which have already examined the genome-wide profile of T-UCRs in other neoplasms like chronic lymphocytic leukemia, colorectal and hepatocellular carcinomas [40], neuroblastoma [41], hepatocellular [42], and prostate [43] carcinomas. The expression profiles obtained from T-UCRs in the current study differed from those observed in previous studies, suggesting that the expression profile of T-UCRs may be tissue and cancer specific, which is reminiscent of tissue-specific expression of miRNAs in human cancers [44].

For the first time, we constructed a three-layered regulatory network ([three component network](https://www.ndexbio.org/#/network/404a7bb1-cc91-11ec-b397-0ac135e8bacf?accesskey=74406b2038d11866bf3aefcc7a7d7946ab61187806f3e198876e1d67d6056891)), containing T-UCRs, miRNAs, and mRNAs specifically for gastric cancer to investigate the complex interactions among these various regulatory RNA levels. In addition, Bioinformatics analyses were performed to reveal the potential roles of T-UCRs in GC. The finally constructed network contains 34 T-UCRs, 275 miRNAs, and 796 mRNAs. As hub nodes are considered as crucial players in the network with respect to the disease-causing gene [45], the T-UCR hubs in the network were defined by nodes with large degree centrality values. Of them, two candidate T-UCRs, uc.232 and uc.343, with an upregulated pattern in TCGA tumoral vs. non-tumoral tissues were selected for further experimental analysis. While uc.232 was undetectable in examined gastric tissues, uc.343 showed an increased expression pattern (although not statistically significant) in tumoral gastric tissues. The latter was consistent with the results obtained from the TCGA cohort. Of note, there are no reports on expression analyses of uc.232 and uc.343 in any tissue/disease, yet. The reason for the discrepancy observed between the results of uc.232 expression patterns in the TCGA cohort vs. our examined tissue samples may be attributable to our small sample size and inherent tumor heterogeneity. It should be noted that in almost 30% of TCGA-STAD, no count was detected for uc.232.

Using GO vocabularies and KEGG signaling pathway information, we analyzed network-related mRNAs. T-UCRs have been demonstrated to regulate mRNA levels by functioning as a sponge for miRNAs, implying that T-UCRs’ biological activity is linked to that of their target mRNAs [39]. Through gene ontology analysis, we discovered that T-UCRs’ target mRNAs are mainly involved in several biological processes, such as regulation of transcription, signal transduction, regulation of cell proliferation, apoptosis, and cell cycle, which are among the most frequently dysregulated processes in cancer [46]. The enriched KEGG pathways included”pathways in cancer”,”miRNAs in cancer”, and”PI3K-AKT signaling pathway”. Cancer cell proliferation, migration, and differentiation have all been found to be mediated by the PI3K-AKT signaling pathway [47].

Protein-protein interaction (PPI) networks offer information on how proteins interact with others to create biological processes within the cell [48, 49]. We identified 20 hub genes in the PPI network based on node degree centralities. MAPK1 and PIK3R1 had the highest degree centralities. Several studies have shown that the MAPK1 gene interacts with miRNAs and lncRNAs to regulate cell proliferation and cell cycle progression in GC [50, 51]. The PI3K pathway is an essential player in tumorigenesis and the ubiquitously hyperactivated signaling pathway in neoplasms [52]. Upregulation of PIK3R1, the regulatory subunit of PI3K, has been shown to cause enhanced cisplatin resistance in GC leading to limited overall clinical efficacy in patients [53]. NEAT1-induced upregulation of PIK3R1 is reported to promote GC progression [54].

A total of five DET-UCRs were revealed as survival-related T-UCRs in STAD using univariate survival analysis. Patients with GC who had higher expression of uc.169 and uc.360 and lower expression of uc.8, uc.342, and uc.417 had considerably shorter survival rates. Our findings are consistent with those obtained by Terreri et al. [55] who reported association of low levels of uc.8+ with a worse overall survival in bladder cancer patients. To the best of our knowledge, none of uc.169, uc.360, uc.342, and uc.417 have been previously studied in neoplasms.

## Conclusions

In this study, we analyzed the expression profiles of T-UCRs in a large number of patients with GC from TCGA cohort, which can serve as a basis for related studies in the field. We furthermore identified dysregulated T-UCRs that might be prospective clinical diagnostic markers and are associated with tumorigenesis and development of GC. A total of five T-UCRs were identified as potential prognostic T-UCRs that were significantly correlated with overall survival. Our study highlights important implications of T-UCRs as novel biomarker candidates for cancer diagnosis and outcome prediction. Thus, our data lays a foundation for further needed functional research of T-UCRs in clinical and pre-clinical GC studies.

## Supplementary files

The data that supports the findings of this study (supplementary files) are available in the Mendeley data repository (https://doi.org/10.17632/ywt9mf6xtn.1).

## Author contributions

Conceptualization, Parvaneh Nikpour; Data curation, Anis Khalafiyan, Markus Wolfien and Ali Salehzadeh-Yazdi; Funding acquisition, Parvaneh Nikpour; Methodology, Anis Khalafiyan, Modjtaba Emadi-Baygi, Markus Wolfien, Ali Salehzadeh-Yazdi and Parvaneh Nikpour; Project administration, Parvaneh Nikpour; Resources, Modjtaba Emadi-Baygi and Parvaneh Nikpour; Supervision, Parvaneh Nikpour; Validation, Parvaneh Nikpour; Writing – original draft, Anis Khalafiyan, Modjtaba Emadi-Baygi, Markus Wolfien, Ali Salehzadeh-Yazdi and Parvaneh Nikpour; Writing – review & editing, Anis Khalafiyan, Modjtaba Emadi-Baygi, Markus Wolfien, Ali Salehzadeh-Yazdi and Parvaneh Nikpour.

## Institutional Review Board Statement

The study was conducted according to the guidelines of the Declaration of Helsinki, and approved by the Ethics Committee of Isfahan University of Medical Sciences (IR.MUI.MED.REC.1398.443).

## Informed Consent Statement

Prior to participating, all patients gave written informed consent to the Iran Tumoral Bank.

## Data Availability Statements

The data that support the findings of this study are available on request from the corresponding author.

## Supplementary files legends

**Supplementary file 1. T-UCR-miRNA and miRNA-mRNA interaction predictions**. The first sheet indicates the predicted interactions between significantly differentially expressed T-UCRs and miRNAs retrieved from RNAhybrid. The second sheet shows the interactions between miRNAs (from the first tab) and mRNAs predicted by miRWalk3.0 online database.

**Supplementary file 2. Functional annotations of network mRNAs**. GO biological processes, molecular functions and KEGG pathways related to 796 mRNAs of the network and retrieved from DAVID are presented in the first, second, and third sheets, respectively. DAVID: The Database for Annotation, Visualization and Integrated Discovery, GO: gene ontology, KEGG: Kyoto Encyclopedia of Genes and Genomes

**Supplementary file 3. KEGG pathways related to the top 20 predominant nodes of the PPI network**. These 20 proteins were considered as an input to STRING online database and then KEGG pathway functional enrichment was retrieved from the analysis tab. KEGG: Kyoto Encyclopedia of Genes and Genomes, PPI: protein-protein interaction

**Supplementary file 4. PCR products on agarose gels**. Electrophoresis of the PCR products on agarose gel revealed single bands with the predicted sizes for the amplified uc.232 and uc.343 segments.

**Supplementary file 5. Sequencing analysis of uc.232 and uc.343 PCR products**. A part of sequence electropherogram of uc.232 (A) and uc.343 (B) and their nucleotide BLAST are presented. BLAST: Basic Local Alignment Search Tool

